# BindPred: A Framework for Predicting Protein-Protein Binding Affinity from Language Model Embeddings

**DOI:** 10.1101/2025.09.27.678407

**Authors:** Haixing Piao, Veda Sheersh Boorla, Somtirtha Santra, Costas D. Maranas

## Abstract

**Motivation:** Reliable predictions of protein–protein binding affinities are essential for molecular biology and therapeutic discovery. However, most computational methods rely on three-dimensional structural models, which are often unavailable for many complexes.

**Results:** We introduce BindPred, a structure-agnostic input framework that predicts affinities directly from amino acid sequences by combining embeddings from large protein language models with gradient boosting trees. On the PPB-Affinity benchmark, which comprises 11,919 diverse complexes, BindPred achieves a Pearson correlation coefficient of 0.86 in random split five-fold cross-validation, where the training and test sets share <30% global sequence identity. Ablation analysis indicates that evolutionary embeddings alone capture most of the predictive signals, while augmenting with physics-based energy terms from PyRosetta and BindCraft increases the correlation by only 0.01. A more stringent protein-level split that places entire protein families (wild-type and all mutants) exclusively in either training or testing sets, results in only a modest decline in performance, demonstrating robust generalization to novel interaction pairs. Because BindPred operates exclusively on sequence input, it enables rapid inference (approximately 3 million complexes per GPU (T4) hour), making proteome-scale screening computationally feasible.

**Availability:** The pretrained model and inference pipeline are available in a Google Colab notebook: <u>BindPred Colab notebook</u>. The training dataset, code, and model weights are available on the Hugging Face: BindPred

**Supplementary information:** Supplementary data are available at *Bioinformatics* online.

## 1 Introduction

Protein–protein interactions (PPIs) govern essential cellular processes from immune recognition to metabolic regulation, making accurate binding affinity prediction (*K*_*d*_ or Δ*G*_*bind*_) indispensable for both fundamental biology and drug discovery. Experimental techniques such as surface plasmon resonance (Sparks et al., 2019) and isothermal titration calorimetry (Csibra and Stan, 2022) provide precise measurements but require purified proteins and specialized equipment, limiting throughput to hundreds rather than the millions of interactions needed for proteome-wide studies.

Computational approaches have emerged along two primary directions. Physics-based methods, including molecular dynamics (MD), free-energy perturbation (FEP), and thermodynamic integration, estimate Δ*G*_*bind*_ by decomposing it into enthalpic and entropic contributions from electrostatics, hydrogen bonding, van der Waals forces, and conformational entropy (Singh and Warshel, 2010; Kříž et al., 2023). These mechanistically rigorous methods require several demanding inputs: high-quality starting structures, carefully tuned force fields, and extensive simulation trajectories to achieve convergence. Even under optimized protocols, calculations often take hours per complex (Liu et al., 2023). Structure-based machine learning (David, Garcia, Reza Haydarlou, & Feenstra, 2025; Chaves et al., 2025; Lebedenko, Polovinkin, Kazovskaia, & Skrynnikov, 2025) accelerates prediction by learning features derived from protein complexes yet remains bottlenecked by structure generation. Even advanced predictors such as AlphaFold2 (Jumper et al., 2021), AlphaFold-Multimer (Evans et al., 2021), and AlphaFold3 (Abramson et al., 2024) frequently misplace binding partners, produce overlapping chains, or distort binding interfaces, especially for multichain assemblies with weak or absent coevolutionary signals (Bryant, Pozzati, and Elofsson, 2022; Wee & Wei, 2024). These limitations have motivated the development of approaches that bypass structure prediction entirely.

Protein language models (pLMs) address this challenge by extracting structural and functional information directly from sequences. Trained on hundreds of millions of sequences, models such as the Evolutionary Scale Model 2 (ESM2) (Lin et al., 2023) and the Multimeric Interaction Transformer (MINT) (Ullanat et al., 2025) learn context-aware embeddings that capture evolutionary constraints, local chemistry, and functional motifs without requiring explicit structural input. These embeddings enable structure-free inference in a single forward pass and have proven effective for function annotation and mutation effect prediction (Lam et al., 2024).

We present BindPred, a structure-free binding affinity predictor integrating frozen pLM embeddings with gradient boosting trees (GBT). Operating directly on sequences, BindPred avoids errors from docking or homology modeling and applies to complexes lacking reliable structures. The embeddings, derived from ESM2 or MINT, remain fixed during training and serve as task-agnostic biochemical descriptors. With only ∼6,000 trainable parameters, the architecture operates on precomputed embeddings and can evaluate approximately three million complexes per GPU (T4) hour, achieving speeds that are orders of magnitude faster than FEP protocols or structure-based machine learning pipelines.

Several models have incorporated pLM embeddings into protein– protein interaction predictors, but most focus on classification (Liu, Young, et al., 2024), relative ΔΔG prediction (Gurusinghe et al., 2024), or interface-specific architectures (Sargsyan & Lim, 2024; Banerjee et al., 2025). ProBASS (Gurusinghe et al., 2024) exemplifies the latter by combining ESM2 embeddings with structure-conditioned vectors from ESM-IF1 (Hsu et al., 2022) to predict ΔΔ*G* upon mutation. Although effective on SKEMPI-style datasets, ProBASS requires high-confidence structural models for each variant and predicts relative changes in affinity, making it unsuitable for ranking entirely novel complexes. In contrast, BindPred directly estimates absolute *log*_10_*K*_*d*_ values from amino acid sequences, eliminating structural dependencies, enabling high-throughput screening, and maintaining strong predictive accuracy across diverse protein interactions.

We evaluated BindPred against a sequence–structure hybrid baseline on the PPB Affinity benchmark (Liu et al., 2023), which spans antibody– antigen, enzyme–inhibitor, and receptor–ligand complexes. Using random split five-fold cross-validation and strict protein-level splits, we assessed predictive accuracy and generalizability. We also trained variants of BindPred that incorporated structural features from PyRosetta (Chaudhury et al., 2010) and BindCraft (Pacesa et al., 2025) to quantify their added value. Across all settings, the embedding-only model matched the augmented variants, achieving a Pearson correlation coefficient of 0.86 under stratified splitting and maintaining strong performance on entirely unseen proteins. Because it operates directly from sequence, BindPred can rapidly screen millions of protein pairs, making it a practical tool for proteome-wide interaction mapping and early-stage therapeutic de novo sequence design.

## 2 Methods

### 2.1 Overview of the BindPred Framework

### 2.2 Dataset and feature extraction

#### 2.2.1 “Corpus PPB-Affinity” dataset

We refer to Corpus PPB-Affinity as the combined dataset of 11,919 protein–protein complexes drawn from SKEMPI v2.0, SAbDab, PDBbind 2020, Affinity Benchmark v5.5, and ATLAS, and distinguish it from the structure-dependent PPB-Affinity baseline model (Liu et al., 2024). For BindPred models that combined embeddings with physical energy terms, we used 9,597 complexes with PyRosetta (19 energy features) and 3,468 complexes with BindCraft (14 energy features), the latter reduced due to segmentation errors.

#### 2.2.2 Sequence feature extractions

Sequence-level embeddings were obtained by mean pooling the residue-level hidden states from the final transformer layer of either the ESM-2 (esm2_t33_650M_UR50D; Lin et al., 2023) model, pretrained on UniRef50 with 650 M parameters, or MINT (Ullanat et al., 2025)(see Supplementary Information for more details).

#### 2.2.3 Structural feature extractions

Structural conformation and stability features were computed using PyRosetta (Chaudhury, Lyskov, and Gray, 2010). All input structures were first relaxed with the FastRelax protocol under the ref2015 score function to remove steric clashes and optimize side-chain packing. Each relaxed structure was then rescored in PyRosetta 4 (release 2024.12) using default ref2015 weights with the *flags ignore_unrecognized_res true, ex1*, and *ex2aro* to enable thorough side-chain sampling. For every wild-type and mutant pose, we extracted the full set of nineteen ref2015 all-atom energy terms, recording both the raw energy values and fourteen derived interaction-energy features. The ref2015 scoring framework is described in detail by Alford et al. (2017), and the interaction-energy calculation protocol follows the BindCraft methodology (Pacesa et al., 2025). The BindCraft energy calculations fail when there are multichain (e.g, two receptor chains with one ligand chain or vice versa), so only two-chained complexes were used for the BindCraft energy calculations.

### 2.3 Cross-validation split strategies and model training

#### 2.3.1 Random split and Protein level fivefold cross-validation

Corpus PPB-Affinity datasets were randomly split (random seed = 42) into training (80%) and testing (20%) sets to optimize the model weights on a subset of data while evaluating its performance on unseen examples. The code can be found on train.py

Protein-level splitting was applied to prevent information leakage by grouping each wild-type complex with all its variants under the same PDB_ID, ensuring that no wild-type–mutation pairs crossed the train–test boundary. Model performance was assessed using five-fold group-aware cross-validation, maintaining an overall 80% training and 20% testing ratio. In each fold, one group served as the test set and the remaining four as the training set.

BindPred was trained using the CatBoost gradient boosting framework with root mean square error (RMSE) as the objective function. Detailed hyperparameters and feature variants are provided in Supplementary Methods.

## 3 Results

We evaluated BindPred across multiple benchmarking scenarios to assess its predictive accuracy and generalization capability. We began by comparing sequence-only models based on ESM2 and MINT embeddings to a hybrid baseline on the comprehensive Corpus PPB-Affinity dataset and its individual subsets. We then performed cross-dataset validation to test robustness under domain shifts, followed by runs using a stringent protein-level data split to assess performance on entirely unseen proteins. Finally, we explored the applicability of BindPred in ranking de novo protein designs, providing a comprehensive assessment of its accuracy and scalability across diverse prediction settings.

Table S1 and Figure 2 provide an overview of the training dataset.

**Figure 1.**
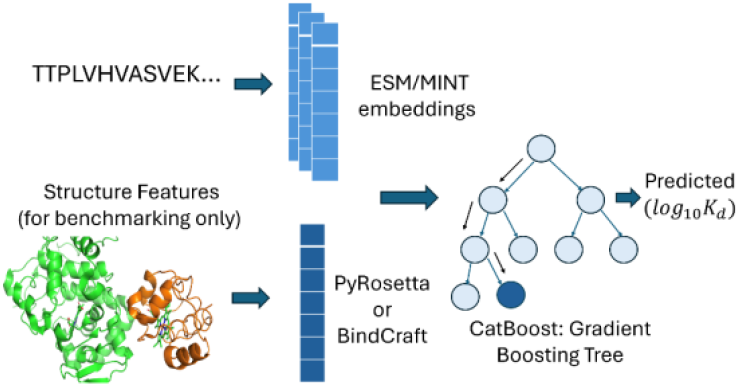
BindPred model architecture and workflow. Protein sequences are encoded with ESM2 or MINT embeddings and used to train a CatBoost regressor to predict *log*_10_*K*_*d*_. Structural energy terms are optionally appended for benchmarking but excluded from the default model. The default model in the Google Colab notebook only requires the ESM2 embedding inputs.

**Figure 2.**
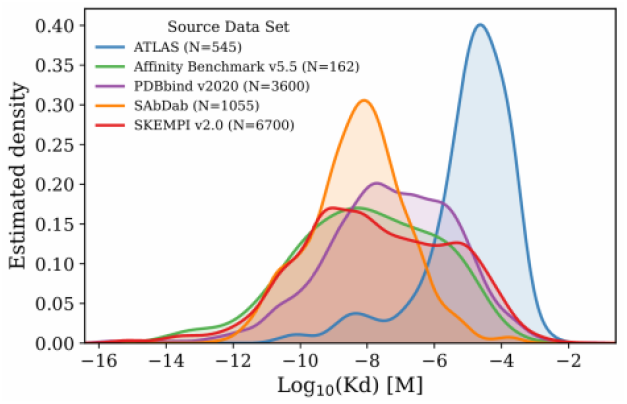
Binding affinity distributions of Corpus PPB-Affinity. Kernel density estimates of *log*_10_*K*_*d*_for protein–protein complexes from each subset. Curves are normalized to unit area to facilitate comparison of distribution shapes (see summary table in the Supplemental Information).

**Figure 3.**
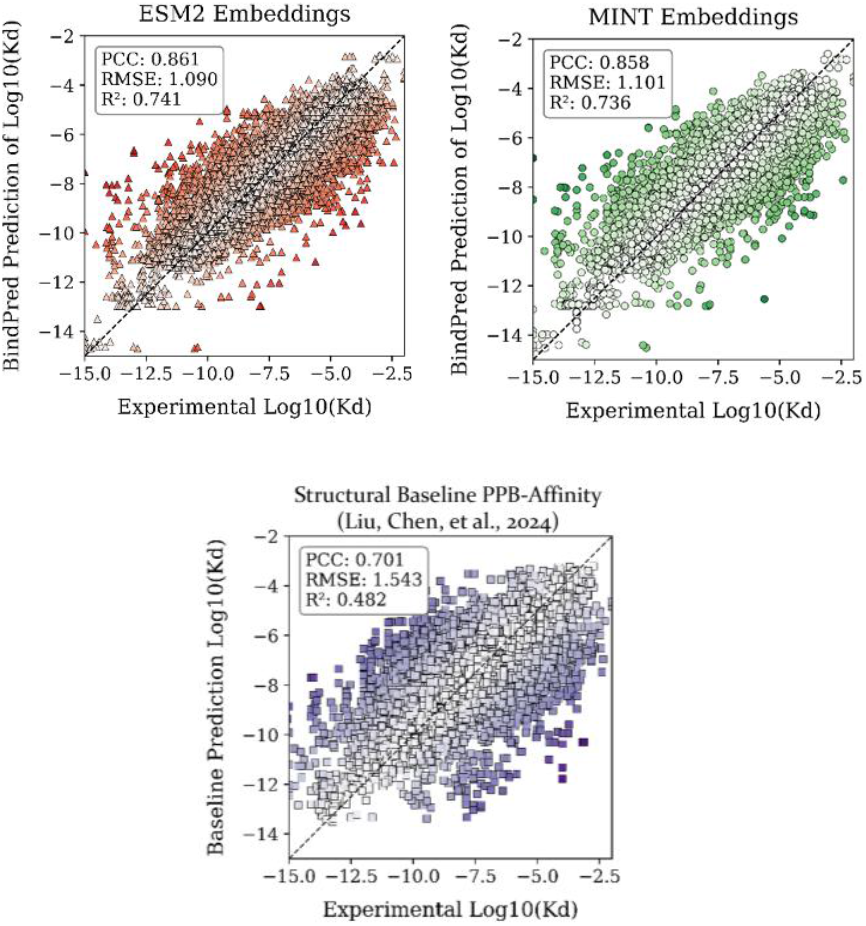
Sequence-only BindPred outperforms the Structural Baseline PPB-Affinity model in random split fivefold cross-validation. Predicted *log*_10_*K*_*d*_versus experimental values for 11,919 protein–protein complexes under random split five-fold cross-validation. Points are colored by absolute error, with closer alignment to the diagonal indicating higher predictive accuracy. BindPred with ESM2 embeddings and with MINT embeddings both outperform the baseline PPB-Affinity model.

**Figure 4.**
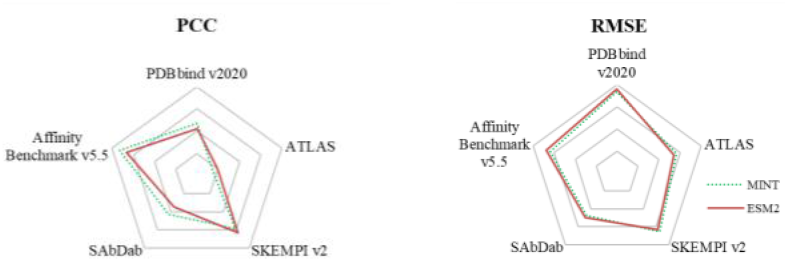
Cross-subset “Stranger” validation. Radar plots show the performance of MINT-based (green, dashed) and ESM2-based (red, solid) BindPred models when trained on four datasets and tested on the fifth. Performance varies substantially by held-out dataset, with Affinity Benchmark v5.5 yielding the highest cross-dataset accuracy and ATLAS showing the largest performance drop. Differences between MINT and ESM2 are modest but consistent, with MINT achieving higher PCC and lower RMSE, suggesting better resilience to domain shifts, although neither model fully overcomes the challenges posed by heterogeneous or structurally distinct interaction types.

**Figure 5.**
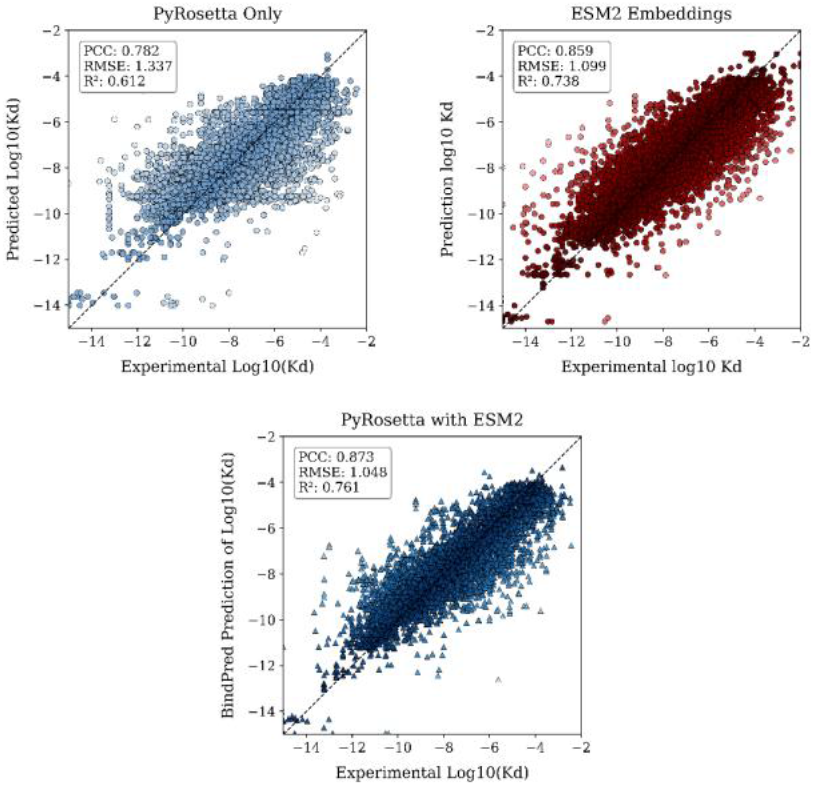
Random split fivefold cross validations across N = 9,597 protein protein complexes for three feature sets. PyRosetta Only uses a 20-dimensional vector of interface energy terms derived from PyRosetta. ESM2 embeddings use 2,560 features obtained by mean pooling receptor and ligand embeddings and concatenating them. PyRosetta with ESM2 combines both for a 2,580-dimensional input. All three variations have been trained using the same protein complexes and hyperparameters indicated in the method section.

**Figure 6.**
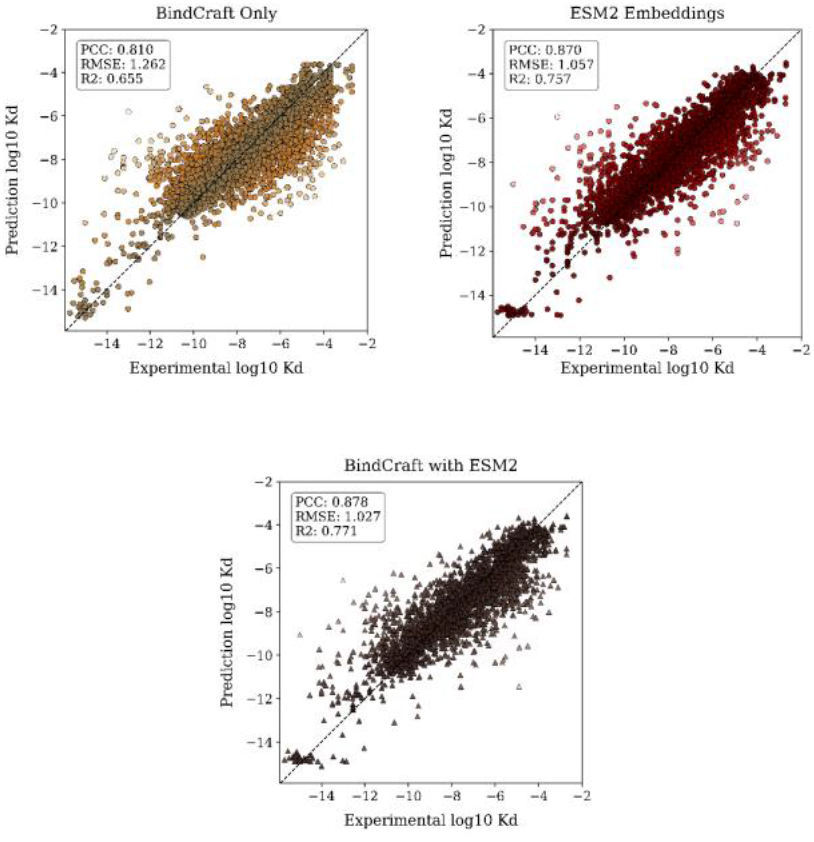
Random split fivefold cross validations across N = 3,917 protein protein complexes for three feature sets. The BindCraft analysis was restricted to 3,468 complexes (∼30% of the benchmark) for which interface energies could be computed successfully. Nearly half of this subset came from SKEMPI v2 entries, which are enriched in point mutations, while the remaining complexes were drawn from four other sources with more heterogeneous interaction types. BindCraft frequently failed for multichain assemblies (e.g., two receptor chains with one ligand or vice versa), so this subset should be viewed as a biased sample rather than a representative slice of the full dataset. To assess the contribution of these features, we trained three models on this subset: ESM2 embeddings only, BindCraft interaction energies only, and a combined model that concatenated both.

**Figure 7.**
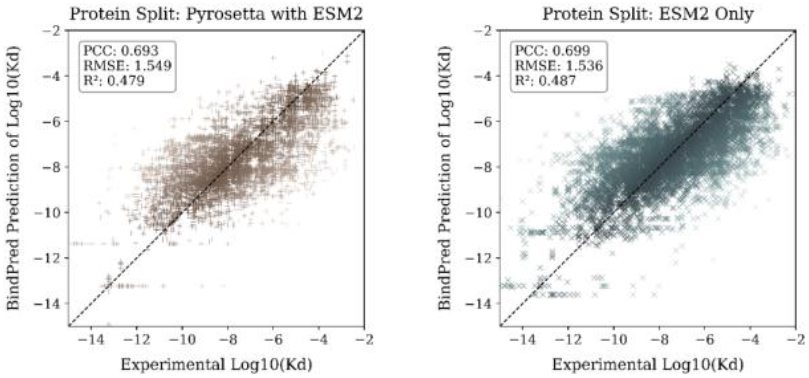
Protein-level split fivefold cross validations. Scatter plots compare predicted versus experimental *log*_10_*K*_*d*_ for protein-level five-fold cross-validation, where all variants of a protein are placed entirely in training or testing sets. Both ESM2 alone and the PyRosetta with the ESM2 model exhibit lower performance under protein split than under random split, as expected from a stricter test of generalization. However, their performance remains closely comparable, even though the latter includes structural energy terms.

### 3.1 Fivefold cross-validation on the Corpus PPB-Affinity

BindPred was evaluated on the full Corpus PPB-Affinity (12,062 total with 11,919 valid complexes) using two protein embedding approaches: ESM2 and MINT, and compared against the PPB-Affinity baseline model. ESM2 provides general-purpose protein embeddings trained on large-scale protein sequence data. MINT extends the 650M-parameter ESM2 architecture with cross-chain attention mechanisms that explicitly model inter-chain residue interactions. MINT undergoes additional training on a curated protein–protein interaction dataset from STRING (Szklarczyk et al., 2024), enabling it to capture interface-specific binding determinants and contextual partner information beyond what single-chain embeddings encode. These design differences suggest that MINT should be advantageous in scenarios where accurate modeling of partner-specific interfaces is important, particularly for complexes with limited evolutionary coupling or atypical binding modes. However, because BindPred uses full-length sequences rather than cropped interface regions, MINT’s interface-specific advantages may be diluted, and in some cases could lead to overfitting if non-interfacial sequence patterns correlate spuriously with binding affinity in the training data.

Under Random Split Five-Fold Cross-Validation in Figure 2, both embedding-based BindPred models exceeded the Structural Baseline PPB-Affinity’s PCC by approximately 0.16. This substantial improvement reflects fundamental differences in how each method represents protein complexes. ESM2- and MINT-based BindPred generate fixed-length 2,560-dimensional embeddings for the full complex by concatenating 1,280-dimensional pooled embeddings for the receptor and ligand chains. Large-scale pretraining endows these representations with information on residue coevolution, long-range dependencies, physicochemical tendencies, and interface propensities directly from sequence. This enables the models to infer binding-relevant features without structural inputs avoiding the resolution variability that can impair structure-dependent pipelines. The PCC difference between ESM2 and MINT was minimal (i.e., 0.003), suggesting that MINT’s interface-aware features did not confer a measurable advantage in the full-sequence evaluation. MINT’s interface-aware features may be diluted by non-interfacial sequence regions, reducing its advantage over ESM2’s general-purpose embeddings.

In contrast, the Structural Baseline PPB-Affinity crops each complex to 128 interface residues and constructs two handcrafted feature sets, dominated by 3D coordinates, by an invariant point attention stack. The per-residue (“node”) features include amino acid identity, sequence position, backbone dihedral angles, and mutation mask embeddings (256-dimensional per residue). The per-pair (“edge”) features include relative positions, inter-residue distances, and geometric orientation encodings (36-dimensional per residue pair). Because every residue is paired with every other residue, the edge representation scales quadratically (O(N^2^)) with the number of residues, producing 128×128=16,384 residue pairs per complex. This increases computational cost and magnifies noise propagation, since an error in a single residue’s coordinates will affect all of its pairwise features. Due to sensitivity to experimental and modeling errors, force-field biases, and variability in structural resolution, these factors can limit generalization when structurally dissimilar complexes appear in the test set.

Despite their overall advantage, the embedding-based models exhibit three limitations. First, extreme affinity outliers are present. For example, the Colicin E2 and Im2 complex (PDB 3U43, experimental *log*_10_*K*_*d*_ = −14.4) was overestimated by ESM2 with a value of −6.2 and MINT at −6.9, with only partial correction from the structural baseline at −7.7. In addition, an engineered TNF alpha affibody (PDB 2TGP, experimental −6.0) was underpredicted by both sequence-only BindPred models at −12.6. Both cases involve deeply buried interfaces near 2,200 Å^2^ and dense salt bridge networks that are not explicitly encoded in sequence embeddings and are only coarsely captured by baseline descriptors. Second, for certain mutations, the embeddings show limited sensitivity because both ESM2 and MINT primarily encode evolutionary statistics from the sequence. When a substitution carries a weak or ambiguous evolutionary signal in the alignment, the models infer little effect and predict that the mutant will behave exactly as the wild type. Third, not surprisingly, performance declines for underrepresented complex families with few close analogs in training. Together, these limitations are pictorially manifested with a broader flat-band effect in which a weak evolutionary signal or lack of close analogs causes predictions to cluster near the mean.

### 3.2 Fivefold cross-validation on individual subset

As mentioned in Figure 1, the Corpus PPB-Affinity dataset is composed of five subsets originating from different source databases, each reflecting distinct experimental conditions, complex types, and structural quality criteria. Evaluating BindPred and the Structural Baseline PPB-Affinity model on each subset separately enables us to assess how well the models generalize across heterogeneous data sources and to identify performance trends that differences in interface size, sequence diversity, or experimental resolution may drive. This breakdown also helps pinpoint subset-specific challenges that may not be apparent in the aggregated results.

The Affinity Benchmark v5.5 subset yielded the highest overall performance for both ESM2 and MINT embedding of BindPred. This benchmark dataset is curated to include high-resolution structures, reliable experimental affinities, and a balanced distribution across the affinity spectrum. Many complexes have deep multiple sequence alignments (MSAs) with clear coevolutionary signal, which pretrained embeddings such as ESM2 and MINT can exploit to encode residue–residue dependencies and partner-specific interface preferences. These conditions allow sequence-only models to accurately capture both the ranking and magnitude of binding affinities.

The SKEMPI v2.0 dataset also exhibits strong correlation performance with BindPred (i.e., PCC = 0.89), reflecting its emphasis on well-characterized complexes with a predominance of single amino acid substitutions. Approximately three-quarters of the entries involve only one residue change in an otherwise identical wild-type background, with most of the remaining cases consisting of double and, more rarely, triple substitutions. This results in a highly controlled mutational landscape in which the global fold, interface geometry, and residue–residue interaction network remain unchanged. Under these conditions, sequence embeddings can effectively capture the full wild-type context while sensitively detecting localized perturbations, as embedding differences can be directly attributed to altered physicochemical properties at or near the substitution site. Consequently, the embedding-based BindPred models can more readily map specific mutations to their effects on binding affinity.

Dataset ATLAS posed the greatest challenge for both ESM2 and MINT embeddings. This dataset is highly heterogeneous, covering a wide range of protein–protein interaction types, many of which are transient or involve substantial conformational changes upon binding. Experimental affinities originate from diverse assay platforms with varying precision, and structural models range from high-quality crystal structures to less certain modeled conformations. Many complexes in ATLAS also lack deep MSAs, especially for rare or artificial constructions, resulting in weak or ambiguous evolutionary signal. Because ESM2 and MINT rely on statistical regularities learned from sequence data, weak evolutionary constraints cause the models to regress toward the mean and lose calibration for absolute binding energy prediction. Nevertheless, for all datasets, BindPred retains significant prediction dominance over the existing state-of-the-art.

### 3.3 Cross-subset validation

The subsets differ substantially in both their interaction types and how binding affinity values were obtained and curated. Affinity Benchmark v5.5 (Guest et al., 2021) compiles affinities primarily from high-quality biophysical experiments, including surface plasmon resonance (SPR) and isothermal titration calorimetry (ITC), with values verified against original publications and converted to binding free energies where needed. SKEMPI v2 (Jankauskaite et al., 2019) derives most of its entries from targeted mutagenesis studies present in the literature, where binding affinity values for wild-type and mutant complexes, measured by SPR, ITC, or enzyme-linked assays, are used to calculate ΔΔ*G*. ATLAS (Borrman et al., 2017) aggregates interaction data from multiple public resources and high-throughput studies, combining *K*_*d*_, *K*_*i*_, and *IC*_50_ measurements from a broad range of assay types, such as fluorescence polarization, AlphaScreen, yeast two-hybrid derivatives, and various biochemical binding assays. By training on four of these subsets and testing on the remaining one (“stranger” validation), we can evaluate how differences in interaction scope, measurement methods, assay precision, and curation standards influence the observed generalization patterns.

Both versions of BindPred have obtained moderate predictive power when trained on sets of interactions and tested on interactions never seen during training, as in the cross-subset stranger validation. PCC values in this setting are lower than in the subset-specific fivefold cross-validation, reflecting the challenges of domain shift, yet both ESM2 and MINT retain meaningful accuracy. MINT embeddings achieve higher PCC in PDBbind v2020 (0.472 vs. 0.423 for ESM2), SAbDab (0.427 vs. 0.346 for ESM2), and the Affinity Benchmark v5.5 set (0.727 vs. 0.667 for ESM2), whereas ESM2 embeddings perform better in ATLAS (0.194 vs. 0.147) and SKEMPI (0.629 vs. 0.587) datasets.

Protein language model MINT was trained exclusively on approximately 96 million protein–protein interaction pairs from the STRING (Szklarczyk et al., 2024) database. STRING interactions are defined from diverse functional evidence, coexpression, gene neighborhood proximity, text mining, and evolutionary co-occurrence rather than high-resolution structural data. While this provides massive sequence-pair diversity, most pairs lack experimentally resolved interfaces. Consequently, pLM MINT’s ligand-aware cross-chain attention learns to approximate interfacial regions indirectly, identifying covariation and conserved sequence motifs that statistically correlate with contact residues across many examples, but without structural confirmation.

When test datasets contain complexes with stable geometries and conserved contact motifs, such as PDBbind v2020 and SAbDab, these STRING-derived interface priors align well with the true binding determinants, enabling MINT embeddings to amplify relevant features and suppress non-interfacial noise. In contrast, ATLAS includes heterogeneous and conformationally dynamic complexes, often with shallow or biased MSAs, where STRING-based patterns may misidentify interface residues or overlook distal contributors to binding. SKEMPI v2’s mutation-centric design further limits MINT’s advantage, as many impactful changes lie outside the interface, making full-sequence modeling from ESM2 embeddings more effective.

### 3.5 Structure-based energy features from PyRosetta and BindCraft

Recognizing the limitations of sequence-only approaches, including flat band effects and failures at extreme binding affinities, structural features were added to the training dataset. ESM2 embeddings were selected for augmentation due to their general-purpose nature, whereas MINT was excluded because its ligand-aware mechanism already encodes partner-specific interfacial context, which may overlap with explicit structural features. This overlap is plausible since MINT’s pretraining may capture much of the interfacial energetic information, reducing the potential benefit of additional structural descriptors. To test the effect of such descriptors, PyRosetta energies from 9,597 complexes capturing van der Waals, solvation, and hydrogen bonding, and BindCraft energies from about complexes quantifying interface-specific energetics via a fast relaxation protocol, were appended to ESM2 embeddings.

Performance improves as the input dimensionality increases from PyRosetta only to ESM2 embeddings, with a smaller but consistent gain when adding the 20 PyRosetta terms to the 2,560-dimensional embedding input, indicating diminishing returns once most of the signal is captured by the embeddings. PyRosetta-only focuses on conformability and stability through packing, hydrogen bonding, and solvation style terms rather than evolutionary sequence context. The embedding model achieves similar accuracy whether trained on 11,919 complexes or on 9,597, suggesting that performance is already near a data sufficiency plateau for this architecture and objective. Adding the 20 PyRosetta energies to the 2,560-dimensional embedding input produces only a small incremental change that does not appear significant relative to fold-to-fold variability. Even though including physical features resolved some flat band issues, this improvement is only noticeable when the mutations cause significant conformational or stability changes. This pattern implies considerable overlap between the information captured by sequence embeddings and the physics descriptors using the PyRosetta energy terms. In practical terms, embedding alone already encodes much of the interfacial energetic context, leaving limited headroom for additional gains.

BindCraft features represent interface-level interaction energies and stability metrics, not explicit pairwise interactions. When the embedding-only model is retrained on the same subset of complexes that successfully produced BindCraft energies, it maintains good performance. Adding the BindCraft feature vector to the 2,560-dimensional embedding input leads to only a small change because the embeddings already capture much of the interfacial energetic context, label noise is lower for this curated subset, and ceiling effects are limited for improvement. In short, the additional features are largely redundant on this dataset. A small number of entries show all zero BindCraft vectors, which arise from interface detection failures, chain or residue misalignment, missing atoms or nonstandard residues that cause the contact graph to be empty, or a silent failure that defaults to zeros. Excluding these edge cases does not alter the overall conclusion that embeddings alone already explain most of the variance, with BindCraft providing limited incremental benefit on this subset.

### 3.6 Protein split

Protein split is essential for a fair test of generalization. In pairwise affinity prediction, many rows are mutants of the same wild type or close homologs, so random splits place near duplicates in both training and testing, letting a model memorize scaffold-specific features, reuse the same alignment signal, and interpolate around the wild type, which could inflate correlation and deflate error. Protein split groups records by a complex identifier such as PDB ID and places the wild type and all mutants entirely in either training or testing, blocking leakage from shared scaffold, interface geometry, and sequence context. Although metrics often drop and fold variance can rise, this evaluation better reflects performance on unseen proteins and provides more trustworthy guidance for design and screening.

This similarity suggests that adding physics-based descriptors contributes little information beyond what the embeddings already encode, and therefore does not materially improve accuracy under a protein split. Protein embeddings are sensitive to residue identity, packing, and the local chemical environment, which explains their strong stand-alone performance. Practically, the embedding-only model achieves comparable accuracy at lower computational cost and complexity. Any incremental benefit from explicit energy features is more likely to appear in “harder” regimes, for example, when the sequence signal is weak, chemistries are unusual, alignment depth is limited, or the evaluation imposes distribution shifts that extend beyond a protein split.

### 3.7 De novo protein sequence design using BindPred

BindPred can rank designed sequences by predicted affinity but does not necessarily correctly classify binding versus non-binding. To test rankings, a small *de novo* design set from the BindCraft supplementary dataset was used. We compared BindPred’s predictions with Rosetta’s Δ*G*_*REU*_, a structure-based energy score commonly used to rank designs (see Supplementary Information for details).

## 4. Discussion

BindPred demonstrates that protein language model embeddings can predict absolute binding affinity with high accuracy, achieving a Pearson correlation coefficient of approximately 0.86 when trained on 11,919 complexes. Incorporating physics-based descriptors from PyRosetta or BindCraft yielded only small gains under fivefold cross-validation. When training and testing proteins were separated by protein identity, both the embedding-only model and the PyRosetta with embedding model declined in performance relative to random splits, as expected, but their results remained closely comparable. This indicates that explicit energy terms provide little incremental information once proteins in the training and testing sets are disjoint.

On the curated BindCraft subset, the embedding-only model already achieved a high correlation, and the addition of interaction energy vectors produced only marginal changes. Notably, this subset was disproportionately composed of SKEMPI v2 mutation entries. The negligible improvement observed is therefore important, as these cases should, in principle, have been most sensitive to interface-specific features. A small number of complexes produced zero BindCraft vectors due to interface detection failures, chain or residue mismatches, missing atoms, or preprocessing errors; excluding these entries did not alter the conclusions.

The comparable performance of gradient boosting and deep neural networks suggests that feature quality may be more important than architectural choice for predicting binding affinity. A simple gradient boosting regressor trained on embeddings matched the accuracy of more elaborate neural networks, consistent with prior studies (Reim et al., 2025; Wu, Chang, & Zou, 2024), which found that improvements from ESM2 embeddings were largely insensitive to model architecture. The limited benefit of explicit energy terms may reflect overlap with information already encoded in embeddings, which capture residue identity, local packing, and chemical environment. The mixed target-wise association between Rosetta Δ*G*_*REU*_ and experimental affinity further underscores the contrast between static structural energetics and sequence-derived evolutionary signals, and cautions against treating Δ*G*_*REU*_ as a proxy for measured binding affinity. In practice, embeddings paired with a simple learner provide a strong and efficient baseline. Future work could explore other embedding approaches or fine-tuning strategies for difficult cases where evolutionary information is limited.

## Acknowledgements

This material is based upon work supported by the Center for Bioenergy Innovation (CBI), U.S. Department of Energy, Office of Science, Biological and Environmental Research Program under Award Number ERKP886. Any opinions, findings, and conclusions or recommendations expressed in this publication are those of the author(s) and do not necessarily reflect the views of the U.S. Department of Energy. This work was also supported by the U.S. National Science Foundation funded Molecule Maker Lab Institute (MMLI), award number 2019897 supported by National AI Research Institutes Program of the Directorate for Computer and Information Science and Engineering (CISE), in collaboration with the Division of Chemistry (CHE) and the Division of Chemical, Bioengineering, and Environmental Transport Systems (CBET) awarded to CDM. The funders had no role in study design, data collection and analysis, decision to publish, or preparation of the manuscript. Computations for this research were performed on the Pennsylvania State University’s Institute for Computational and Data Sciences’ Roar supercomputer.

---

**Table 1.** **Five-fold cross-validation results for each individual subset. Benchmark = Affinity Benchmark v5.5; PPB = Structural Baseline PPB-Affinity.**

| Subset | PCC |  |  | RMSE |  |  |
| --- | --- | --- | --- | --- | --- | --- |
|  | ESM2 | MINT | PPB | ESM2 | MINT | PPB |
| **Benchmark** | 0.895 | 0.886 | 0.671 | 1.011 | 1.052 | 1.618 |
| **ATLAS** | 0.632 | 0.657 | 0.387 | 0.963 | 0.934 | 1.169 |
| **PDBbind** | 0.824 | 0.820 | 0.662 | 1.175 | 1.188 | 1.627 |
| **SAbDab** | 0.661 | 0.666 | 0.335 | 0.954 | 0.948 | 1.447 |
| **SKEMPI** | 0.898 | 0.891 | 0.745 | 0.891 | 0.916 | 1.544 |

**Table 2.**
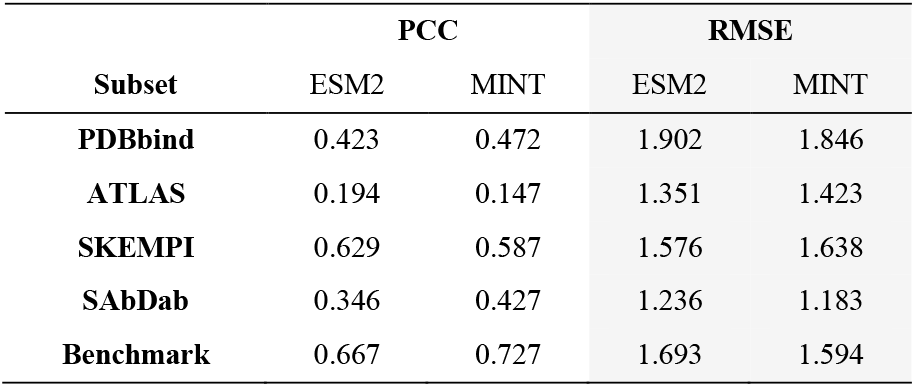
Cross-subset stranger validation metrics.

## References

Abramson, J., Adler, J., Dunger, J., Evans, R., Green, T., Pritzel, A., … Beattie, C. (2024). Accurate structure prediction of biomolecular interactions with AlphaFold 3. Nature, 630(630), 493–500. doi: 10.1038/s41586-024-07487-w

Alford, R. F., Leaver-Fay, A., Jeliazkov, J. R., O’Meara, M. J., DiMaio, F. P., Park, H., … Gray, J. J. (2017). The Rosetta All-Atom Energy Function for Macromolecular Modeling and Design. Journal of Chemical Theory and Computation, 13(6), 3031–3048. doi: 10.1021/acs.jctc.7b00125

Banerjee, S., Karimi, M., Yilmaz, M., Jaakkola, T., Dubrov, B., Shang, S., & Benson, R. (2020). Pi-SAGE: Permutation-invariant surface-aware graph encoder for binding affinity prediction. Retrieved August 11, 2025, from Arxiv.org website: https://arxiv.org/html/2508.01924v1

Bateman, A., Martin, M.-J., Orchard, S., Magrane, M., Ahmad, S., Alpi, E., … Insana, G. (2023). UniProt: the Universal Protein Knowledgebase in 2023. Nucleic Acids Research, 51(D1). doi: 10.1093/nar/gkac1052

Borrman, T., Cimons, J., Cosiano, M., Purcaro, M., Pierce, B. G., Baker, B. M., & Weng, Z. (2017). ATLAS: A database linking binding affinities with structures for wild-type and mutant TCR-pMHC complexes. Proteins: Structure, Function, and Bioinformatics, 85(5), 908–916. doi: 10.1002/prot.25260

Bryant, P., Pozzati, G., & Elofsson, A. (2022). Improved prediction of protein-protein interactions using AlphaFold2. Nature Communications, 13(1), 1265. doi: 10.1038/s41467-022-28865-w

Chaudhury, S., Lyskov, S., & Gray, J. J. (2010). PyRosetta: a script-based interface for implementing molecular modeling algorithms using Rosetta. Bioinformatics, 26(5), 689–691. doi: 10.1093/bioinformatics/btq007

Chaves, E., Sartori, J., Santos, W. M., Cruz, C., Mhrous, E. N., Nacimento-Filho, M. F., … Lins, R. D. (2025). Estimating Absolute Protein–Protein Binding Free Energies by a Super Learner Model. Journal of Chemical Information and Modeling. doi: 10.1021/acs.jcim.4c01641

Csibra, E., & Stan, G.-B. (2022). Absolute protein quantification using fluorescence measurements with FPCountR. Nature Communications, 13(1). doi: 10.1038/s41467-022-34232-6

David Garcia, C.M., Reza Haydarlou, & Feenstra, K. A. (2025). PIPENN-EMB ensemble net and protein embeddings generalise protein interface prediction beyond homology. Scientific Reports, 15(1). doi: 10.1038/s41598-025-88445-y

Eddy, S. R. (2011). Accelerated Profile HMM Searches. PLoS Computational Biology, 7(10). doi: 10.1371/journal.pcbi.1002195

Evans, R., O’Neill, M., Pritzel, A., Antropova, N., Senior, A., Green, T., … Kohli, P. (2021). Protein complex prediction with AlphaFold-Multimer. BioRxiv. doi: 10.1101/2021.10.04.463034

Guest, J. D., Thom Vreven, Zhou, J., Moal, I. H., Jeliazkov, J. R., Gray, J. J., … Pierce, B. G. (2021). An expanded benchmark for antibody-antigen docking and affinity prediction reveals insights into antibody recognition determinants. Structure, 29(6), 606-621.e5. doi: 10.1016/j.str.2021.01.005

Gurusinghe, S. N. S., Wu, Y., DeGrado, W., & Shifman, J. M. (2024). ProBASS – a language model with sequence and structural features for predicting the effect of mutations on binding affinity. doi: 10.1101/2024.06.21.600041

Hsu, C., Verkuil, R., Liu, J., Lin, Z., Hie, B., Sercu, T., … Rives, A. (2022). Learning inverse folding from millions of predicted structures. doi: 10.1101/2022.04.10.487779

Jankauskaite, J., Jiménez-García, B., Justas Dapkūnas, Fernández-Recio, J., & Moal, I. H. (2019). SKEMPI 2.0: an updated benchmark of changes in protein–protein binding energy, kinetics and thermodynamics upon mutation. Bioinformatics, 35(3), 462–469. doi: 10.1093/bioinformatics/bty635

Jumper, J., Evans, R., Pritzel, A., Green, T., Figurnov, M., Ronneberger, O., … Back, T. (2021). Highly Accurate Protein Structure Prediction with Alphafold. Nature, 596(7873), 583–589. doi: 10.1038/s41586-021-03819-2

Lam, H. Y. I., Guan, J. S., Ong, X. E., Pincket, R., & Mu, Y. (2024). Protein language models are performant in structure-free virtual screening. Briefings in Bioinformatics, 25(6). doi: 10.1093/bib/bbae480

Lebedenko, O. O., Polovinkin, M. S., Kazovskaia, A. A., & Skrynnikov, N. R. (2025). PCANN Program for Structure-Based Prediction of Protein–Protein Binding Affinity: Comparison With Other Neural-Network Predictors. Proteins: Structure, Function, and Bioinformatics, 93(9), 1498–1506. doi: 10.1002/prot.26821

Lin, Z., Akin, H., Rao, R., Hie, B., Zhu, Z., Lu, W., … Rives, A. (2023). Evolutionary-scale prediction of atomic-level protein structure with a language model. Science, 379(6637), 1123–1130. doi: 10.1126/science.ade2574

Liu, D., Young, F., Lamb, K. D., Claudio Quiros, A., Pancheva, A., Miller, C., … Yuan, K. (2024). PLM-interact: extending protein language models to predict protein-protein interactions. doi: 10.1101/2024.11.05.622169

Liu, H., Chen, P., Zhai, X., Huo, K.-G., Zhou, S., Han, L., & Fan, G. (2024). PPB-Affinity: Protein-Protein Binding Affinity dataset for AI-based protein drug discovery. Scientific Data, 11(1). doi: 10.1038/s41597-024-03997-4

Marks, D. S., Colwell, L. J., Sheridan, R., Hopf, T. A., Pagnani, A., Zecchina, R., & Sander, C. (2011). Protein 3D Structure Computed from Evolutionary Sequence Variation. PLoS ONE, 6(12), e28766. doi: 10.1371/journal.pone.0028766

Marquet, C., Heinzinger, M., Olenyi, T., Dallago, C., Erckert, K., Bernhofer, M., … Rost, B. (2021). Embeddings from protein language models predict conservation and variant effects. 141(10), 1629–1647. doi: 10.1007/s00439-021-02411-y

Morcos, F., Pagnani, A., Lunt, B., Bertolino, A., Marks, D. S., Sander, C., … Weigt, M. (2011). Direct-coupling analysis of residue coevolution captures native contacts across many protein families. Proceedings of the National Academy of Sciences, 108(49). doi: 10.1073/pnas.1111471108

Pacesa, M., Nickel, L., Schellhaas, C., Schmidt, J., Ekaterina Pyatova, Kissling, L., … Goverde, C. A. (2025). One-shot design of functional protein binders with BindCraft. Nature. doi: 10.1038/s41586-025-09429-6

Pavel Kříž, Beránek, J., & Vojtěch Spiwok. (2023). Free Energy Differences from Molecular Simulations: Exact Confidence Intervals from Transition Counts. Journal of Chemical Theory and Computation, 19(7), 2102–2108. doi: 10.1021/acs.jctc.2c01237

Reim, T., Hartebrodt, A., Blumenthal, D. B., Bernett, J., & List, M. (2025). Deep learning models for unbiased sequence-based PPI prediction plateau at an accuracy of 0.65. Bioinformatics, 41(Supplement_1), i590–i598. doi: 10.1093/bioinformatics/btaf192

Remmert, M., Biegert, A., Hauser, A., & Söding, J. (2012). HHblits: lightning-fast iterative protein sequence searching by HMM-HMM alignment. Nature Methods, 9(2), 173–175. doi: 10.1038/nmeth.1818

Sargsyan, K., & Lim, C. (2024). Using protein language models for protein interaction hot spot prediction with limited data. BMC Bioinformatics, 25(1). doi: 10.1186/s12859-024-05737-2

Singh, N., & Warshel, A. (2010). A comprehensive examination of the contributions to the binding entropy of protein-ligand complexes. Proteins: Structure, Function, and Bioinformatics, 78(7), 1724–1735. doi: 10.1002/prot.22689

Sledzieski, S., Singh, R., Cowen, L., & Berger, B. (2021). D-SCRIPT translates genome to phenome with sequence-based, structure-aware, genome-scale predictions of protein-protein interactions. Cell Systems, 12(10), 969-982.e6. doi: 10.1016/j.cels.2021.08.010

Sparks, R. P., Jenkins, J. L., & Fratti, R. (2019). Use of Surface Plasmon Resonance (SPR) to Determine Binding Affinities and Kinetic Parameters Between Components Important in Fusion Machinery. Methods in Molecular Biology (Clifton, N.J.), 1860, 199–210. doi: 10.1007/978-1-4939-8760-3_12

Steinegger, M., & Söding, J. (2017). MMseqs2 enables sensitive protein sequence searching for the analysis of massive data sets. Nature Biotechnology, 35(11), 1026–1028. doi: 10.1038/nbt.3988

Steinegger, M., & Söding, J. (2018). Clustering huge protein sequence sets in linear time. Nature Communications, 9(1), 1–8. doi: 10.1038/s41467-018-04964-5

Suzek, B. E., Wang, Y., Huang, H., McGarvey, P. B., & Wu, C. H. (2015). UniRef clusters: a comprehensive and scalable alternative for improving sequence similarity searches. Bioinformatics, 31(6), 926–932. doi: 10.1093/bioinformatics/btu739

Szklarczyk, D., Nastou, K., Koutrouli, M., Kirsch, R., Mehryary, F., Hachilif, R., … von Mering, C. (2024). The STRING database in 2025: protein networks with directionality of regulation. Nucleic Acids Research, 53(D1). doi: 10.1093/nar/gkae1113

Ullanat, V., Jing, B., Sledzieski, S., & Berger, B. (2025). Learning the language of protein-protein interactions. doi: 10.1101/2025.03.09.642188

Wee, J., & Wei, G.-W. (2024). Benchmarking AlphaFold3’s protein-protein complex accuracy and machine learning prediction reliability for binding free energy changes upon mutation. ArXiv, arXiv:2406.03979v1. Retrieved from https://pubmed.ncbi.nlm.nih.gov/38883239/

Wu, K. E., Chang, H., & Zou, J. (2024). ProteinCLIP: enhancing protein language models with natural language. BioRxiv (Cold Spring Harbor Laboratory). doi: 10.1101/2024.05.14.594226

