## Supplementary for "BindPred: A Framework for Predicting Protein-Protein Binding Affinity from Language Model Embeddings"

### Encoding coevolutionary signals in embeddings

Protein language models, including ESM2 and MINT, rely on evolutionary patterns captured in multiple sequence alignments (MSAs). The depth and diversity of these alignments directly determine the strength of coevolutionary signals (Marks et al., 2011; Morcos et al., 2011). MSAs are generated by searching chain sequences against databases such as UniRef (Suzek, Wang, Huang, McGarvey, & Wu, 2015), UniProt (The UniProt Consortium, 2023), and BFD (Steinegger & Söding, 2018) using alignment tools including HHblits (Remmert, Biegert, Hauser, & Söding, 2012), JackHMMER (Eddy, 2011), and MMseqs2 (Steinegger & Söding, 2017). The resulting alignments stack homologous sequences with equivalent residues in corresponding columns. Across evolutionary time, residues in physical contact or contributing jointly to function undergo correlated substitutions to maintain stability and binding specificity (Morcos et al., 2011). These coevolutionary patterns appear as statistical dependencies between columns, with signal strength correlating with alignment quality. Deep, diverse alignments yield robust signals while shallow datasets produce weak patterns. pLMs such as ESM2 (Lin et al., 2022) and MINT (Ullanat, Jing, Sledzieski, & Berger, 2023) implicitly learn these correlations during pretraining, enabling inference of residue–residue contacts, interface constraints, and partner-specific preferences without structural input. This creates a clear performance divide: complexes with deep, diverse alignments yield accurate predictions, while those with shallow alignments produce weak signals and poor sensitivity to binding determinants.

### Sequence feature extractions

Sequence-level embeddings were obtained by mean pooling the residue-level hidden states from the final transformer layer of either the ESM-2 (esm2_t33_650M_UR50D; Lin et al., 2023) model, pretrained on UniRef50 with 650 M parameters, or MINT (Ullanat et al., 2025), which employs attention-based mechanisms to capture cross-chain residue interactions for accurate prediction of protein–protein interfaces directly from sequence data. For each complex, two amino acid sequences were defined: a receptor $s^R$ of length $L_R$ and a ligand $s^L$ of length $L_L$. If a complex contained more than two chains, chains were grouped into receptor and ligand, respectively, according to biological assembly annotations in the PDB files. Redundant or identical chains (e.g., repeated homodimer subunits) were removed so that each unique sequence was represented once, preserving correct stoichiometry without over-representing repeated chains.

The <cls> token is used by the model to produce a pooled representation summarizing the entire sequence, while the <eos> token marks sequence termination and helps the model learn context boundaries. After the ESM-2 tokenizer prepends <cls> and appends <eos>, the tokenized length of the partner $X \in \{R, L\}$ becomes $L_X + 2$, where $L_X$ is the original sequence length. The ESM2 model maps each tokenized sequence to a hidden state matrix

$$H^{(X)} \in \mathbb{R}^{(L_X+2)\times d}$$

where $d$ is the embedding dimension (d=1,280 for the 650 M-parameter variant). The residue rows (indices 1 through $L_X$) are mean-pooled to obtain a fixed-length vector

$$\tilde{h}^{(X)} = \frac{1}{L_X}\sum_{i=1}^{L_X} H_i^{(X)} \in \mathbb{R}^d,$$

excluding the special token rows. Then, concatenating the two-partner embedding gives the complex level representation

$$v = \left[\tilde{h}^{(R)}; \tilde{h}^{(L)}\right] \in \mathbb{R}^{2d},$$

which are fed into CatBoost to predict the $log_{10}K_d$.

### BindPred training

For implementation, we relied on the CatBoost (https://catboost.ai) framework to train the gradient boosting tree model for predicting binding affinity. The hyperparameters of the model were set at 2,000 iterations, a learning rate of 0.08, and a depth of 4 for all models. The loss function used was the Root Mean Square Error (RMSE), which penalizes larger errors more strongly due to the squaring of residuals and is defined mathematically as:

$$RMSE = \sqrt{\frac{1}{n}\sum_{i=1}^{n}(y_i - \hat{y}_i)^2}$$

Where n is the size of the data point while training, $y_i$ is the actual binding affinity and $\hat{y}_i$ is the predicted binding affinity. The model inputs consisted of ESM2 embeddings, MINT embeddings, ESM2 combined with PyRosetta-derived energy terms, or ESM2 combined with BindCraft energy terms.

Table S1. Summary of the Corpus PPB-Affinity

| Dataset | Primary Interaction Classes | Notes |
|---|---|---|
| SKEMPI v2 | TCR–pMHC, antibody–antigen, protease–inhibitor, other PPIs (mutants) | ~7,085 mutation entries, ~345 wild-type structures |
| PDBbind v2020 | Enzyme–inhibitor, signaling complexes, other natural PPIs | ~2,852 protein–protein binding entries, large dataset but includes mixed labels; some entries may be protein–ligand rather than protein–protein. |
| ATLAS | TCR–peptide–MHC (immune interactions) | ~694 samples, ~112 with crystal structures |
| SAbDab | Antibody–antigen | ~736 entries with both affinity data with structures |
| Affinity Benchmark v5.5 | Antibody, enzyme, signaling PPIs (curated benchmark) | ~207 structures with affinities; high-quality, manually curated benchmark for affinity prediction |

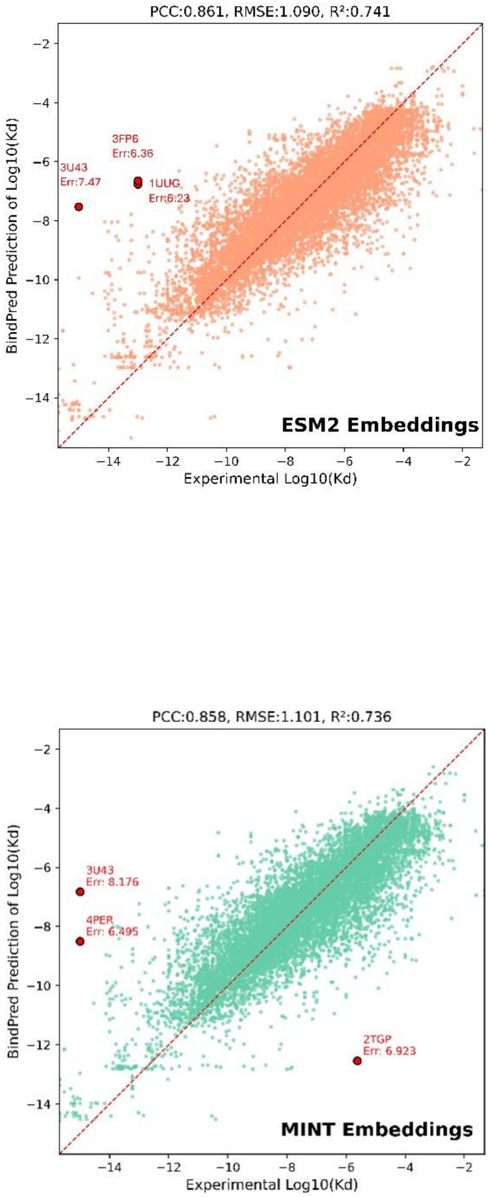


**Figure S1. Comparison of BindPred performance using ESM2 and MINT embeddings on the Corpus PPB-Affinity dataset.** Predicted versus experimental binding affinities are shown, with the three largest error cases highlighted.

### De novo protein sequence design using BindPred

BindPred can rank designed sequences by predicted affinity, but does not necessarily correctly classify binding versus non-binding. To test rankings, a small *de novo* design set from the BindCraft supplementary dataset was used. Many existing workflows rank designs using Rosetta's $\Delta G_{REU}$, a structure-based score in relative energy units derived from packing, hydrogen bonding, and shape complementarity after relaxation. While effective within Rosetta-generated designs, it is important to stress that $\Delta G_{REU}$ omits entropy and solvent effects, is sensitive to model quality, and is not calibrated to experimental units. Relating Rosetta $\Delta G_{REU}$ to experimental affinity is informative because it tests whether a structure-based energy score can serve as a useful proxy for binding strength. $\Delta G_{REU}$ is an internal, uncalibrated estimate of interaction favorability derived from static structures accounting for packing, hydrogen bonding, and solvation. Stronger binding should produce more favorable modeled interactions and therefore more negative $\Delta G_{REU}$. Experimental affinity increases as $K_d$ decreases, and $log_{10}K_d$ becomes smaller for stronger binders. A negative correlation is therefore expected when $\Delta G_{REU}$ is compared with $log_{10}K_d$. If instead $\Delta G_{REU}$ is compared with experimental $\Delta G_{bind}$, a positive association is expected once both quantities are expressed with the same sign convention, since stronger binding corresponds to more negative values for both

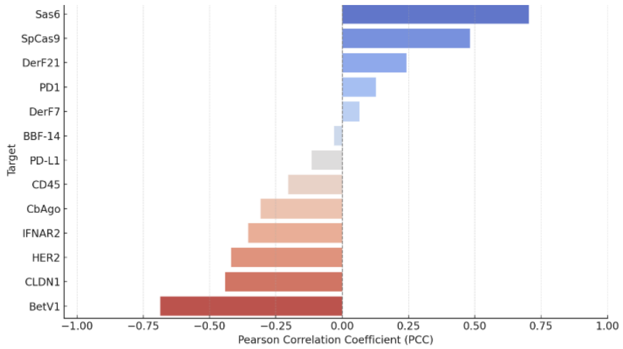


**Figure S2. Correlation between BindPred predictions and Rosetta $\Delta G_{REU}$ for de novo design targets.** Bars show Pearson correlation coefficients (PCC) between BindPred-predicted affinities and Rosetta estimated binding free energies $\Delta G_{REU}$. A negative correlation is expected, as lower Rosetta $\Delta G_{REU}$ typically reflects favorable packing, hydrogen bonding, and shape complementarity.

While some targets follow this trend, others show weak or positive correlations. These differences stem from Rosetta $\Delta G_{REU}$ emphasizing enthalpic terms and omitting entropy and solvent effects, its dependence on structure quality, and the fundamental contrast between BindPred's sequence-based evolutionary signals and Rosetta's static structure energetics. When $K_d$ value is below 1 M, $log_{10}K_d$ is negative and becomes more negative as binding strengthens. Rosetta $\Delta G_{REU}$ likewise becomes more negative when modeled interactions are more favorable. A good design, therefore, exhibits both a low $\Delta G_{REU}$ and a low $log_{10}K_d$. Under these sign conventions, the two quantities should correlate positively across designs because weaker binders move both $\Delta G_{REU}$ and $log_{10}K_d$ toward less negative values, while stronger binders move both toward more negative values. Transforming affinity to $pK_d$ defined as -$log_{10}K_d$ inverts the relation, yielding the expected negative association between $\Delta G_{REU}$ and $pK_d$. Figure 8 shows that combining BindPred with Rosetta $\Delta G_{REU}$ can help prioritize de novo designs, as BindPred captures sequence-driven affinity signals while Rosetta evaluates structural stability.
